# Telemetry validation of dorsal scute microchemistry: complementary tools reconstruct the migratory life-history of an endangered population of Atlantic Sturgeon

**DOI:** 10.1101/2025.01.21.634119

**Authors:** Evan C. Ingram, Matthew E. Altenritter, Kylee B. Wilson, Michael G. Frisk, Kellie R. McCartin

**Author notes:** Deceased (Matthew E. Altenritter).

## Abstract

An understanding of age-specific habitat requirements and the timing of critical ontogenetic transitions is essential to inform conservation and recovery efforts for Atlantic Sturgeon (*Acipenser oxyrinchus*). However, these are often difficult to assess due to the species’ cryptic nature and the consequent lack of temporal and spatial habitat data across life stages. Sampling and microchemistry analysis of dorsal scute apical spines (i.e., dorsal scutes) represents an innovative methodology to reconstruct past life-history events of endangered populations of Atlantic Sturgeon. Here, we establish the broad potential of dorsal scute sampling for wild-caught Atlantic Sturgeon and compare age-estimates and trace-element ratio patterns from dorsal scute with those from pectoral fin spines. We also evaluate microchemistry signatures from both structures to infer past habitat use and age at initial entry into marine waters and, importantly, validate these interpretations using known locations from acoustic telemetry detections. Major ontogenetic shifts in habitat use detected in the microchemistry signatures suggest our methodology was appropriate to identify the timing of initial juvenile migration into marine habitat, while providing additional information regarding the timing of transitions between freshwater and marine habitats that are essential to the management and conservation of this endangered species. As such, the collection of dorsal scute samples is suggested to complement ongoing research efforts for wild-populations and provides additional data points beyond those available from conventional tag-recapture methods alone, allowing researchers and managers to retrospectively identify environmental transitions and habitat use that occur prior to sampling encounters or outside of monitored areas.

## Introduction

The Atlantic Sturgeon (*Acipenser oxyrinchus oxyrinchus*) is an imperiled fish species that exhibits obligate, stage-specific transitions between freshwater spawning habitat and the sea (Bemis and Kynard, 1997; McDowall, 1987; Auer, 1996; Bain, 1997; McDowall, 1997). As such, Atlantic Sturgeon are reliant on access to and the availability of a diversity of functionable freshwater, estuarine, and marine habitats for the successful completion of their life cycle. This specialized anadromous migratory strategy provides spawning populations of Atlantic Sturgeon ecological flexibility across a broad latitudinal range of coastal rivers, from the Saint Lawrence River in New Brunswick to the Saint Marys River in Florida-Georgia (Hilton et al., 2016). This provides individuals the opportunity to exploit conditions and resources that are favorable for growth and reproductive success across aquatic biomes (e.g., sanctuary from marine predators for early life stages in freshwater biomes [Dodson et al., 2009]; faster and larger growth in marine biomes because of access to favorable temperatures and high ecological productivity, [Gross et al., 1988]; potential mechanism for dispersal and rapid recolonization [McDowall, 1987]).

While there are clear adaptive advantages associated with an anadromous life history, the overall complexity of these migratory behaviors has largely confounded management efforts for Atlantic Sturgeon. In the United States, human activities during the 19^th^ and 20^th^ centuries (e.g., unregulated harvest coupled with large-scale degradation of riverine habitat) led to precipitous declines in Atlantic Sturgeon stocks and localized extirpations (Smith 1985; Brumbach 1986; Waldman et al., 1996; Smith and Clugston 1997; Waldman and Wirgin, 1998; Bain et al., 2000; Secor et al., 2002; Daniels et al., 2005; ASSRT, 2007). A moratorium on all Atlantic Sturgeon fisheries in state waters was enacted in 1998 (ASMFC, 1998) and the species was listed under the US Endangered Species Act (ESA) in 2012. The coastwide population of Atlantic Sturgeon in the US is currently designated as distinct population segments (DPS) that are threatened (i.e., Gulf of Maine DPS) or endangered (i.e, New York Bight, Chesapeake Bay, Carolina, and South Atlantic DPSs; USOFR, 2012). Despite these restrictive federal protections, Atlantic Sturgeon stocks remain depleted relative to historical levels (see ASMFC, 2017). Recovery efforts have shown limited results and are further constrained by the unique life-history traits of Atlantic Sturgeon (e.g., long-lived, late-maturing, and anadromous) and obligatory interactions with developed coastal landscapes throughout their life cycle (reviewed in Hilton et al., 2016). Atlantic Sturgeon are still vulnerable as bycatch in commercial fisheries along the Atlantic Coast (e.g., Stein et al., 2004; Beardsall et al., 2013; Dunton et al., 2015) and continue to face emerging threats in coastal systems (Collins et al., 2000; Lenhardt et al., 2006; ASSRT, 2007).

An understanding of age-specific habitat requirements and the timing of critical ontogenetic transitions is essential to informing conservation and recovery efforts for Atlantic Sturgeon. However, these are often difficult to assess due to the species’ cryptic nature and the consequent lack of temporal and spatial habitat data across life stages. While the overall sequence of anadromous behaviors undertaken by Atlantic Sturgeon is relatively predictable across the species’ range (see Hilton et al., 2016), the timing and frequency of specific migratory characters are not well defined and are expected to vary among populations in response to clinal gradients in growth and longevity (e.g., Van Eenennaam and Doroshov, 1998; Stevenson and Secor, 1999; Balazik et al., 2010; Dadswell et al., 2016). Generally, Atlantic Sturgeon are confined to their natal river system for the first two to six years of life (e.g., Dovel and Berggren 1983; Kieffer and Kynard, 1993; Bain 1997; Fox et al., 2018; Fox and Peterson, 2019). Once hatched, larval fish migrate downriver from freshwater spawning habitat into increasingly saline waters before settling into estuarine habitat (e.g., Bain, 1997; Kynard and Horgan, 2002). During the early juvenile stage [i.e., fork length (FL) < 500 mm, based on coastwide NMFS permitting definitions], individuals reside primarily in mesohaline waters below the head of tide before transitioning to a fully marine environment. Marine residents [i.e., late juveniles (FL = 500–1000 mm) and sub-adults (FL = 1000–1300 mm)], undertake extensive seasonal migrations along the coast (e.g., Melnychuk et al., 2017; Ingram et al., 2019; Rothermel et al., 2020), exploiting foraging and growth opportunities and occasionally entering non-natal estuaries (e.g., Savoy and Pacileo, 2003; Savoy, 2007; Kieffer and Kynard, 1993). Atlantic Sturgeon reach sexual maturity at an advanced age (i.e., 7–27 years for females and 5–24 years for males, depending on latitude; Hilton et al., 2016) and will return as adults (FL > 1300 mm) to natal rivers to spawn above the point of salt-incursion every one to five years (e.g., Smith, 1985; Bain, 1997) over an extended lifespan that can last upwards of 60 years (see Hilton et al., 2016).

Research approaches commonly used to study environmental transitions in other fish species have limited efficacy at identifying environmental transitions for Atlantic Sturgeon across the species full life history (see Nelson et al., 2013). Much of the current state of knowledge regarding Atlantic Sturgeon movements and habitat use is the result of extensive surveys and mark-recapture tagging efforts (e.g., Dovel and Berggren 1983; VanEenennaam et al., 1996; Bain 1997; Bain et al., 2000; Peterson et al., 2000; Sweka et al., 2007), as well as genetics analysis (e.g., Waldman et al., 1996; Laney et al., 2007) and—more recently—biotelemetry tracking (e.g., Ingram and Peterson, 2016; Melnychuk et al., 2017; Whitmore and Litvak, 2018; Ingram et al., 2019; Breece et al., 2021). These studies provided the foundational framework for describing the life-stages and seasonal distributions—either in granular fashion relative to salinity, river features, and upstream distances (e.g., Dovel and Berggren 1983; Bain 1997) or through fine-scale description of specific sites of interest (e.g., Sweka et al., 2007). However, these methods are only capable of providing information on habitat use at the time of capture or during subsequent encounters, and do not provide an indication of habitat use prior to sampling. Furthermore, they can be difficult to implement across multiple habitats for long-lived, widely ranging species and are inevitably restricted in scope by practical limitations to spatial or temporal coverage (Heupel and Webber, 2012; Pritt and Frimpong, 2014). This can confound management efforts for rare or endangered species through underestimation of habitat use at intermediate scales or across difficult to monitor areas and is particularly problematic for Atlantic Sturgeon—which undertake extensive, repeated migrations across multiple habitats over an extended lifespan.

The recent development of hard-part microchemistry for fisheries applications has facilitated identification of the environmental history of individual fishes (see Pracheil et al., 2014). Ontogenetic changes in trace element ratios observed in calcified structures such as otoliths and bone tissues have proven useful for retrospectively tracing critical environmental transitions of across salinity gradients (e.g., Secor et al., 1995; Secor and Piccoli, 1996; Veinott et al., 1999; Allen et al., 2009; Hale and Swearer, 2008; Feutry et al., 2012; Hughes et al., 2014; Warburton et al., 2018; Górski et al., 2018). Specific elements can act as useful tracers of environmental history. For example, strontium (SR) in waters with higher salinities is incorporated into calcified structures more efficiently than barium (Ba), which is generally inhibited by salinity (e.g., Tian et al., 2021). As such, differences in elemental ratios of Sr to Calcium (Ca) are generally interpreted as changes in ambient salinity, while changes in Ba to Ca ratios are interpreted as habitat shifts in freshwater (Secor et al., 1995; Kraus and Secor 2004; Walther and Limburg, 2012; Turner and Limburg, 2014). However, the collection of structures commonly used for analysis of trace element ratios can be harmful and/or invasive—particularly in the case of otoliths, which can only be obtained at the expense of sacrificing the sampled individual—and is not generally appropriate when sampling endangered or rare populations in the wild (e.g., Kahn and Mohead, 2010; Baremore and Rosati, 2014; Chalupnicki and Dittman, 2016). Furthermore, the validation of microchemistry inferences from these structures is difficult outside of a laboratory or hatchery setting (where an individual’s age and history of environmental exposure are known) and to our knowledge has not been accomplished for wild populations (see Sellheim et al., 2017).

Dorsal scute apical spines (i.e., dorsal scute) represent a novel sampling structure for the reconstruction of past life history events in Atlantic Sturgeon. Dorsal scutes are modified ganoid scales that are retained throughout an individual’s lifetime and may display chronological layering and a chemical record of past environmental exposure (Magnin, 1963; Thieren and Van Neer, 2016). Importantly, the collection of dorsal scutes is an alternative, non-lethal sampling option that is unlikely to compromise an individual’s swimming performance or health. Altenritter et al. (2015) provided proof of concept that these structures can be used for the reconstruction of past habitat use for the sympatric shortnose sturgeon (*Acipenser brevirostrum*). However, no studies have explored the potential of dorsal scutes for application to microchemistry techniques in Atlantic Sturgeon.

The overall goal of this study was to establish the dorsal scute as a viable structure for the analysis and reconstruction of past life history events in Atlantic Sturgeon. Unlike in laboratory or hatchery settings, Atlantic Sturgeon of known ages are not readily available when working with wild populations. Therefore, endorsement of dorsal scutes as unbiased time-keeping structures capable of reconstructing past exposure to environmental conditions is largely contingent on comparative assessment alongside the pectoral fin spine—a calcified structure that has previously been used to estimate ages of Atlantic Sturgeon (Stevenson and Secor, 1999). As such, our objectives were stepwise, with the initial aim to assess congruence between age estimates and trace-element ratio patterns in a dorsal scute and pectoral fin spine sampled from the same individuals. We then evaluated the microchemistry signatures from these structures to infer past habitat use and age at initial ocean entry. Finally, we validated interpretations of past environmental exposure using known locations obtained from individual acoustic telemetry detection histories.

## Methodology

### Capture and sample processing

All methods for the capture and handling of Atlantic Sturgeon in this study were authorized by the National Marine Fisheries Service (NMFS; Endangered Species Permit 20351), New York State Department of Environmental Conservation (NYSDEC; Endangered/Threatened Species Scientific License 336), and Stony Brook University’s Institutional Animal Care and Use Committee (IRB-1022451–4). Atlantic Sturgeon were captured during research tows at marine aggregation sites off the Rockaway Peninsula, New York (Figure 1; see Ingram et al., 2019). The collection of calcified structures was prioritized for fish that had been tagged with acoustic transmitters during previous sampling efforts. Such individuals had the potential to be detected on fixed arrays of acoustic telemetry receivers maintained throughout the lower Hudson River, New York, and surrounding coastal waters (Figure 1; see Melnychuk et al., 2017; Ingram et al., 2019; Breece et al., 2021). Because receiver arrays were sited throughout a variety of freshwater (< 0.5 ppt), seasonally brackish (0.5–18 ppt), and brackish (18–30 ppt), and offshore marine habitats, telemetry detection histories from tagged fish could be used to ground-truth and validate inferences from microchemistry analysis of hard-parts over a range of environments (Figures 1 and 2).

**Figure 1.**
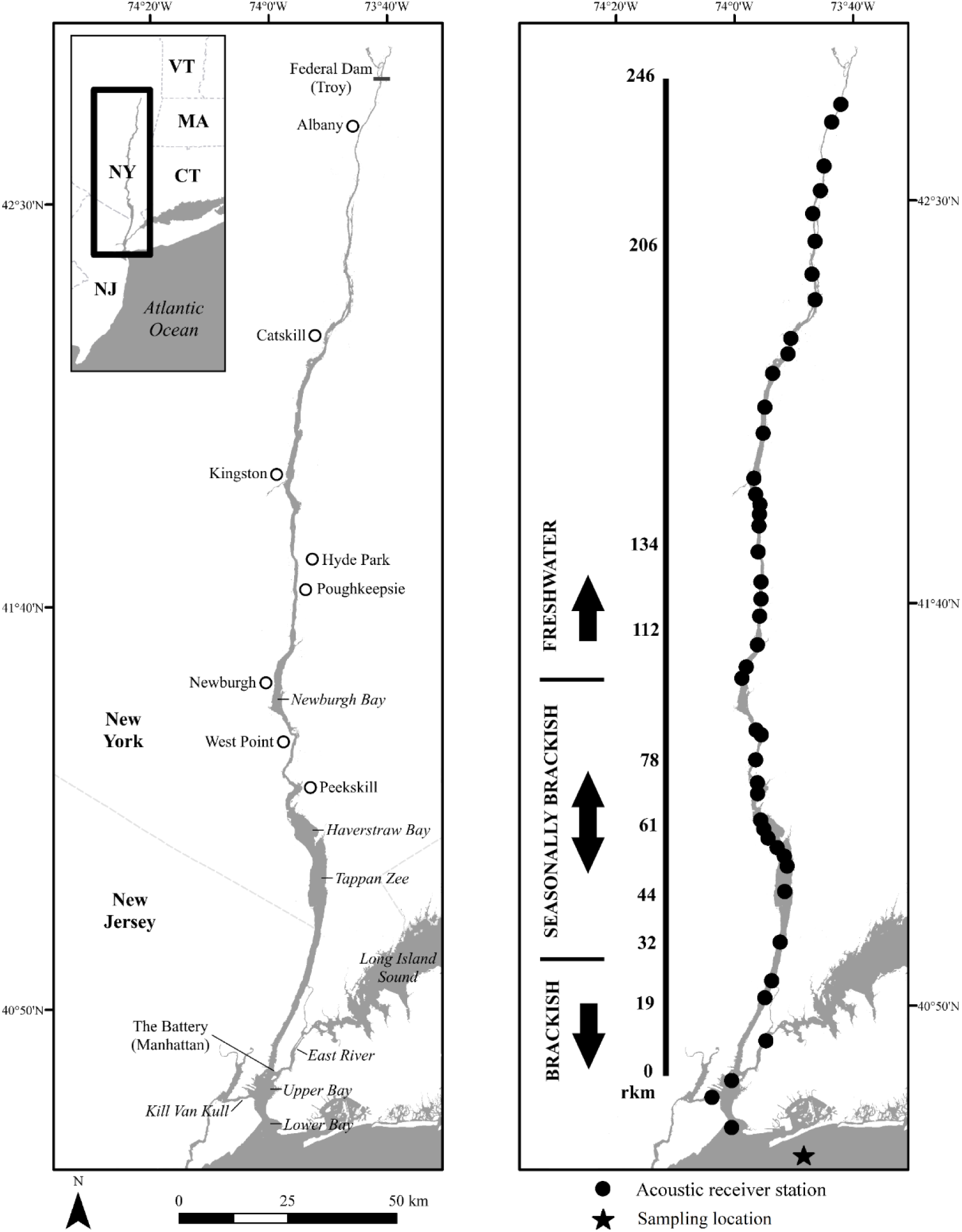
Map of the lower Hudson River in New York and associated water bodies. Geographic features of interest (left panel) and locations of in-river acoustic receiver stations and marine sampling (right panel) are indicated. General river kilometer (rkm) locations of environmental transitions. waters are based on rkm distance above the Battery at Manhattan Island in New York (i.e., rkm 0). Freshwater < 0.5 ppt; Seasonally Brackish = 0.5–18 ppt; Brackish = 18–30 ppt.

**Figure 2.**
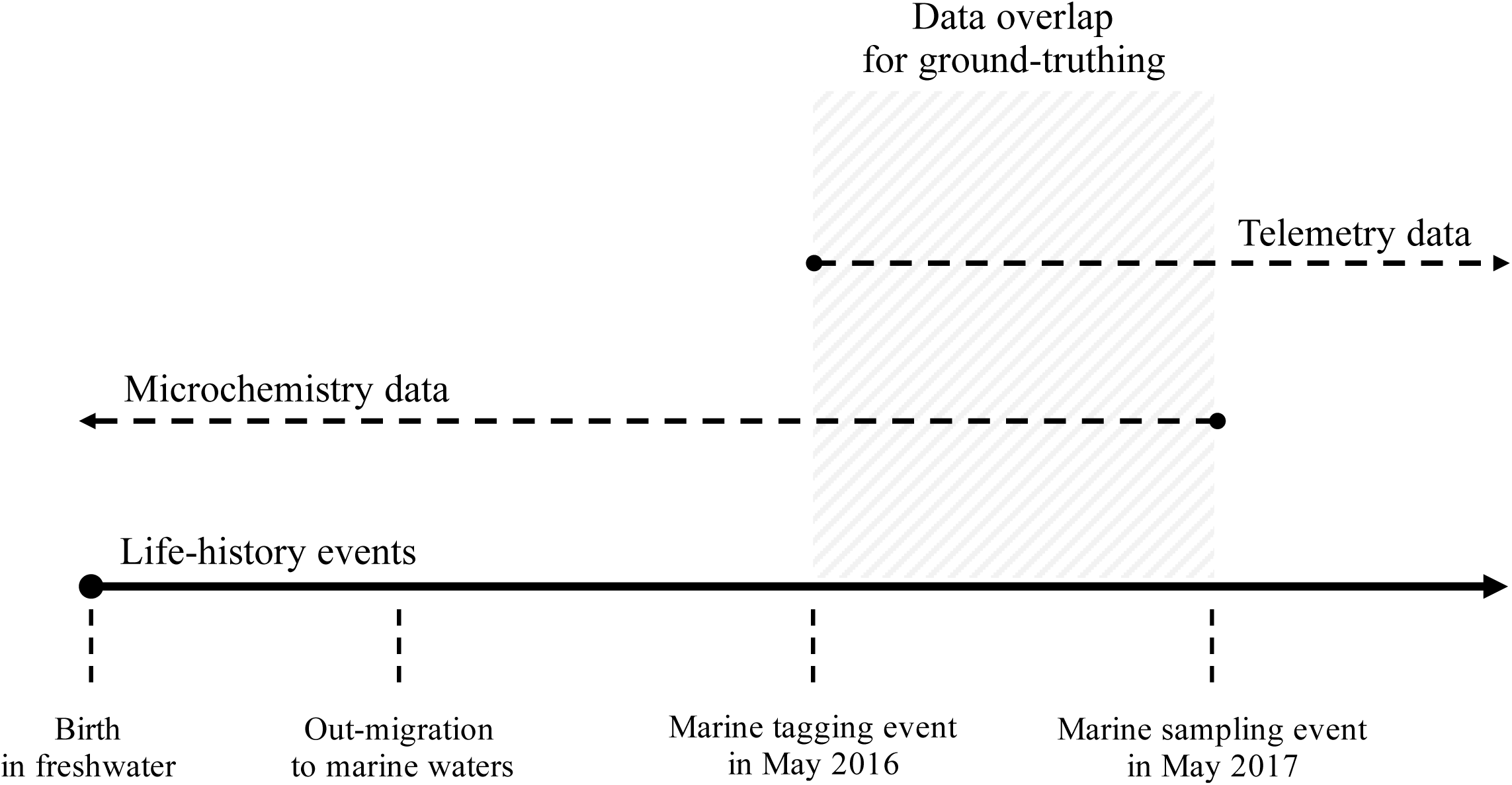
Timeline depicting the use of complementary acoustic telemetry and microchemistry data sources to reconstruct the lifetime history of habitat use for an Atlantic Sturgeon (USFWS ID - 355160038; see Table 1). The period of data overlap between the initial acoustic telemetry tagging event and subsequent recapture and hard-part sampling was used to validate interpretations of microchemistry habitat-use along a salinity gradient. Note, timeline not drawn to scale.

**Table 1.**
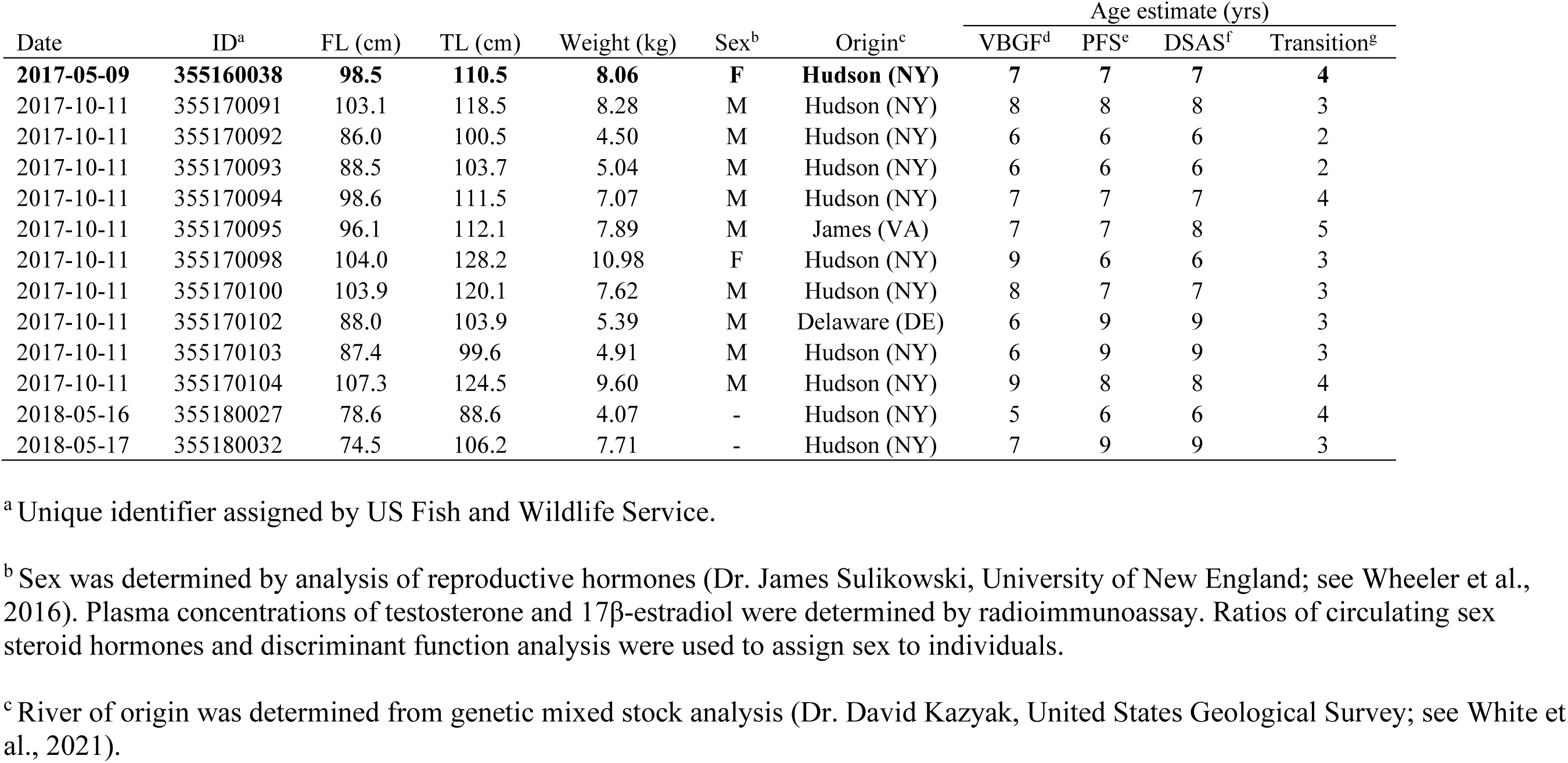
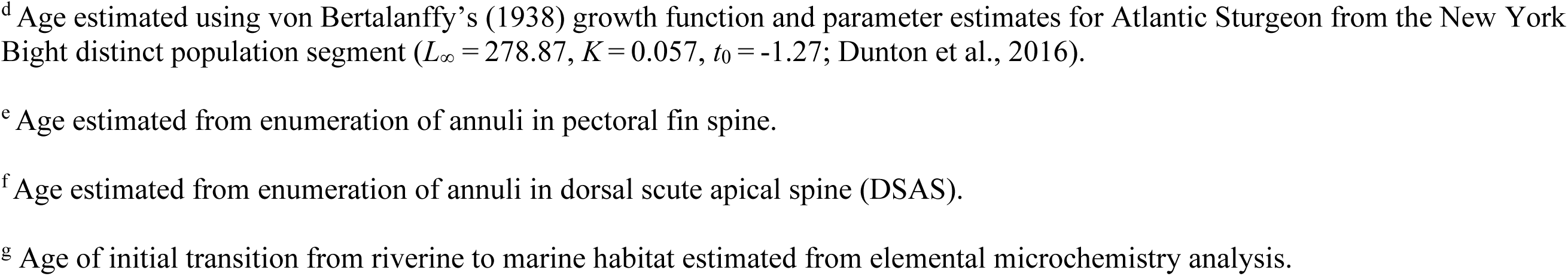
Biological data from Atlantic Sturgeon (n = 13) sampled in marine waters off the Rockaway Peninsula, New York, during 2017 and 2018. Age estimates are provided for comparison, as well as the estimated age of initial transition from riverine to marine waters. Text in bold indicates a recaptured individual that had previously been tagged with an acoustic telemetry transmitter, providing data overlap for the validation of microchemistry inferences. [FL = fork length; TL = total length; VBGF = von Bertalanffy growth function; PFS = pectoral fin spine; DSAS = dorsal scute apical spine].

| Date | ID<sup>a</sup> | FL (cm) | TL (cm) | Weight (kg) | Sex<sup>b</sup> | Origin<sup>c</sup> | Age estimate (yrs) |  |  |  |
| --- | --- | --- | --- | --- | --- | --- | --- | --- | --- | --- |
|  |  |  |  |  |  |  | VBGF<sup>d</sup> | PFS<sup>e</sup> | DSAS<sup>f</sup> | Transition<sup>g</sup> |
| **2017-05-09** | **355160038** | **98.5** | **110.5** | **8.06** | **F** | **Hudson (NY)** | **7** | **7** | **7** | **4** |
| 2017-10-11 | 355170091 | 103.1 | 118.5 | 8.28 | M | Hudson (NY) | 8 | 8 | 8 | 3 |
| 2017-10-11 | 355170092 | 86.0 | 100.5 | 4.50 | M | Hudson (NY) | 6 | 6 | 6 | 2 |
| 2017-10-11 | 355170093 | 88.5 | 103.7 | 5.04 | M | Hudson (NY) | 6 | 6 | 6 | 2 |
| 2017-10-11 | 355170094 | 98.6 | 111.5 | 7.07 | M | Hudson (NY) | 7 | 7 | 7 | 4 |
| 2017-10-11 | 355170095 | 96.1 | 112.1 | 7.89 | M | James (VA) | 7 | 7 | 8 | 5 |
| 2017-10-11 | 355170098 | 104.0 | 128.2 | 10.98 | F | Hudson (NY) | 9 | 6 | 6 | 3 |
| 2017-10-11 | 355170100 | 103.9 | 120.1 | 7.62 | M | Hudson (NY) | 8 | 7 | 7 | 3 |
| 2017-10-11 | 355170102 | 88.0 | 103.9 | 5.39 | M | Delaware (DE) | 6 | 9 | 9 | 3 |
| 2017-10-11 | 355170103 | 87.4 | 99.6 | 4.91 | M | Hudson (NY) | 6 | 9 | 9 | 3 |
| 2017-10-11 | 355170104 | 107.3 | 124.5 | 9.60 | M | Hudson (NY) | 9 | 8 | 8 | 4 |
| 2018-05-16 | 355180027 | 78.6 | 88.6 | 4.07 | - | Hudson (NY) | 5 | 6 | 6 | 4 |
| 2018-05-17 | 355180032 | 74.5 | 106.2 | 7.71 | - | Hudson (NY) | 7 | 9 | 9 | 3 |

<sup>a</sup> Unique identifier assigned by US Fish and Wildlife Service.
<sup>b</sup> Sex was determined by analysis of reproductive hormones (Dr. James Sulikowski, University of New England; see Wheeler et al., 2016). Plasma concentrations of testosterone and 17 $\beta$ -estradiol were determined by radioimmunoassay. Ratios of circulating sex steroid hormones and discriminant function analysis were used to assign sex to individuals.
<sup>c</sup> River of origin was determined from genetic mixed stock analysis (Dr. David Kazyak, United States Geological Survey; see White et al., 2021).
<sup>d</sup> Age estimated using von Bertalanffy's (1938) growth function and parameter estimates for Atlantic Sturgeon from the New York Bight distinct population segment ( $L_{\infty} = 278.87$ , $K = 0.057$ , $t_0 = -1.27$ ; Dunton et al., 2016).
<sup>e</sup> Age estimated from enumeration of annuli in pectoral fin spine.
<sup>f</sup> Age estimated from enumeration of annuli in dorsal scute apical spine (DSAS).
<sup>g</sup> Age of initial transition from riverine to marine habitat estimated from elemental microchemistry analysis.

Paired dorsal scute and pectoral fin spine samples were collected from individual fish. Methodology for dorsal scute sampling was established and refined on hatchery fish at the US Fish and Wildlife Service Northeast Fishery Center (Lamar, PA). Using a fine-toothed hacksaw, a 2–3 cm, wedge-shaped sample of the apical spine was removed from the most-intact dorsal scute by making two shallow-angled cuts perpendicular to the long axis of the fish (Figures S1 and S2, Supporting Information). Similarly, cutting pliers were used to remove a 1–2 cm section of the primary pectoral fin spine at the point of articulation (see Moser et al., 2000; Stevenson and Secor 2000). Samples were air-dried and stored in fish scale envelopes until being processed, allowing soft tissue to undergo microbial decay. Any remaining tissue was removed manually. Samples were then embedded in epoxy and sectioned for subsequent aging and microchemistry analysis (see Appendix S1).

### Age estimation

Sectioned hard-part samples were aged according to established methods to determine correspondence in growth zone counts (Brennan and Calliet, 1989; Stevenson and Secor, 1999). Dorsal scutes and pectoral fin spines were viewed under a microscope using transmitted light to enumerate growth zones. Putative annuli (i.e., growth zones) were defined as pairs of alternating dark (i.e., viewed as opaque under transmitted light) and white bands (i.e., translucent under transmitted light), indicating periods of fast, summer growth or slow, winter growth, respectively (Baremore and Rosati, 2014). Each sample was aged by three experienced readers, without reference to collection date, fish size, or previous estimates. Care was taken to identify and exclude false annuli from annulus counts, with false annuli identified as broken or discontinuous bands that did not circumscribe the entirety of the fin-spine section (e.g., Prince et al., 1985; Stevenson and Secor, 1999). In instances where a secondary (i.e., marginal) fin spine was embedded within the primary fin spine, annuli were only enumerated for the primary spine (e.g., Feindeis, 1997; Stevenson and Secor, 1999).

### Microchemical analysis

Elemental concentrations of ^43^Ca, ^88^Sr, and ^138^Ba in dorsal scute and pectoral fin spine sections were measured using laser ablation inductively coupled plasma mass spectrometry (LA-ICP-MS; State University of New York College of Environmental Science and Forestry). Glass standards (National Institute of Standards and Technology 610) were sampled at the beginning of every session and each sample was preceded by a 30s gas blank to collect background element intensities. Digitized transect paths for dorsal scute sections tracked perpendicularly to observed growth zones from the ventral to the dorsal edge of the structure, while transect paths for pectoral fin sections traversed the entire width of the structure, perpendicular to the orientation of presumed zone formation from the natal core to the posterior edge. Microanalytical carbonate (MACS-3; United States Geological Survey [USGS]) and microanalytical carbonate phosphate standards (MAPS-4; USGS) were analyzed before each sample for calibration purposes and to correct for instrument drift. Transect runs were preceded by a pre-ablation pass (100-μm per s; 110-μm spot size; 10% energy; 0.9-J/cm^2^ fluence) to decontaminate the surface and followed by a 30s gas blank to collect background element intensities. Each transect was ablated with beam settings of 3-μm s^-1^ scan speed and 20% energy and spot size of 85-μm for dorsal scute sections and 35-μm for pectoral fin spines sections. The duration of acquisition for each sample varied based on transect length. Generally, pectoral fin spine transects were around three-times larger than those of dorsal scute, resulting in longer acquisition times.

### Life history transitions and validation

Elemental concentrations of Sr and Ba from microchemistry analysis (reported as element:Ca x 10^3^ ratios) were for plotted against distance from the core for each transect and used as proxies for habitat transition. Changes in Sr:Ca were interpreted as changes in ambient salinity and changes in Ba:Ca were interpreted as habitat shifts in freshwater (Secor et al., 1995; Kraus and Secor 2004; Turner and Limburg, 2014). To standardize the definition of habitat shifts, a regime shift algorithm (Rodionov, 2004) was applied to elemental composition data for Sr:Ca to infer the timing of environmental transitions (e.g., Turner and Limburg, 2012; Altenritter et al., 2015). This algorithm used sequential *t*-tests to identify differences in Sr:Ca values over time, with a moving average used to identify a habitat shift if more than one point was significantly different from the proceeding value (*p* < 0.05). The habitat shift was validated if the difference in means was maintained upon inclusion of subsequent data points (cut off = 10), at which point a new moving average was applied. Available telemetry detection data was used to identify elemental concentration thresholds for Sr:Ca signatures during periods of environmental transition. Environmental data overlap between the initial acoustic telemetry tagging event and subsequent recapture and hard-part sampling was then used to validate microchemistry habitat-use interpretations. This allowed reconstruction of habitat use across the individual’s full life history along a salinity gradient, as elemental signatures of Sr:Ca and Ba:Ca from microchemistry provide information of past environmental use (i.e., from the time of birth until sampling), and telemetry data could be collected from the time of tagging until the expiration of the transmitter’s battery (Figure 2).

## Results

### Fish sampling

Atlantic Sturgeon described in this study (n = 13) were sampled in coastal marine waters of New York during 2016 and 2017 (Figure 1). All fish were identified as late juveniles or sub-adults (i.e., non-mature marine migrants) at the time of capture, based on field measurements of fork-length (FL) and literature definitions (Bain, 1997). The size range of sampled Atlantic Sturgeon was 74.5 to 107.3 cm FL, with a mean FL of 93.4 cm (SD = 10.36; Table 1). Ages-at-capture estimated using the von Bertalanffy growth function (VBGF; von Bertalanffy, 1938) and parameter estimates for Atlantic Sturgeon from the New York Bight distinct population segment (*L*_∞_ = 278.87, *K* = 0.057, *t*_0_ = -1.27; Dunton et al., 2016) ranged from five to nine years (Table 1). Analysis of blood plasma reproductive hormones identified two females and nine males, with the remainder undetermined (Wheeler et al., 2016; Table 1). Most Atlantic Sturgeon were identified to rivers in the New York Bight DPS (total fish = 12; Hudson River = 11 fish; Delaware River = 1 fish), although a single fish originated from the James River in Virginia in the Chesapeake Bay DPS (Table 1).

### Age estimation

Sectioned dorsal scutes and pectoral fin spines displayed distinct growth band patterns (Figure 3). The structure of the dorsal scute apical spine appeared dual-layered, with a thin, mirrored zone located above alternating opaque and translucent zones that were assumed to be annuli (Altenritter et al., 2015; Figure 3). Age estimates from dorsal scutes ranged from six to nine years (mean [SD] = 7.4 [1.2] years) and were generally consistent between paired structures from the same individuals (Table 1). Regression analysis indicated a significant and strong correlation between the number of growth zones found in dorsal scutes and the number of growth zones in pectoral fin spines from the same individual (n = 13; r^2^ = 0.95, *p* < 0.05; Figure 4). The slope of this relationship was not significantly different from unity (*p* = 0.336), suggesting that dorsal scutes can be used in place of pectoral fin spines for aging applications.

**Figure 3.**
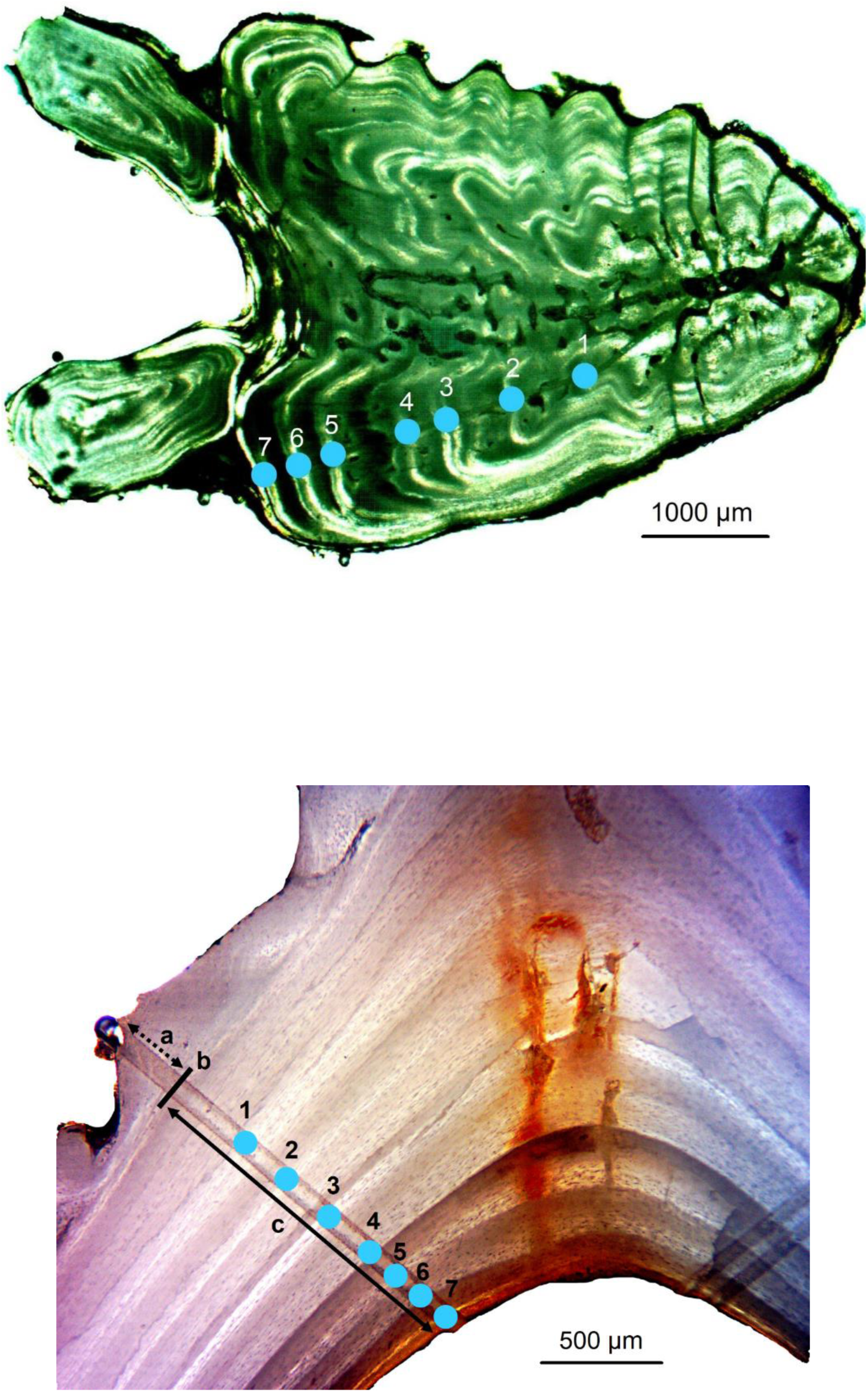
Sectioned and polished pectoral fin spine (top image) and dorsal scute apical spine (bottom image) samples from a wild-caught Atlantic Sturgeon as viewed under transmitted light (shown for USFWS ID - 355160038; see Table 1). The structure of the dorsal scute apical spine is dual-layered and composed of a thin, mirrored zone (a) above alternating opaque and translucent bands (c), separated at the natal boundary (b). Growth zones (single blue circles) are enumerated in both structures and assumed annuli. The estimated age of this individual from both structures is comparable at 7+ years. Note, images are shown at different scales.

**Figure 4.**
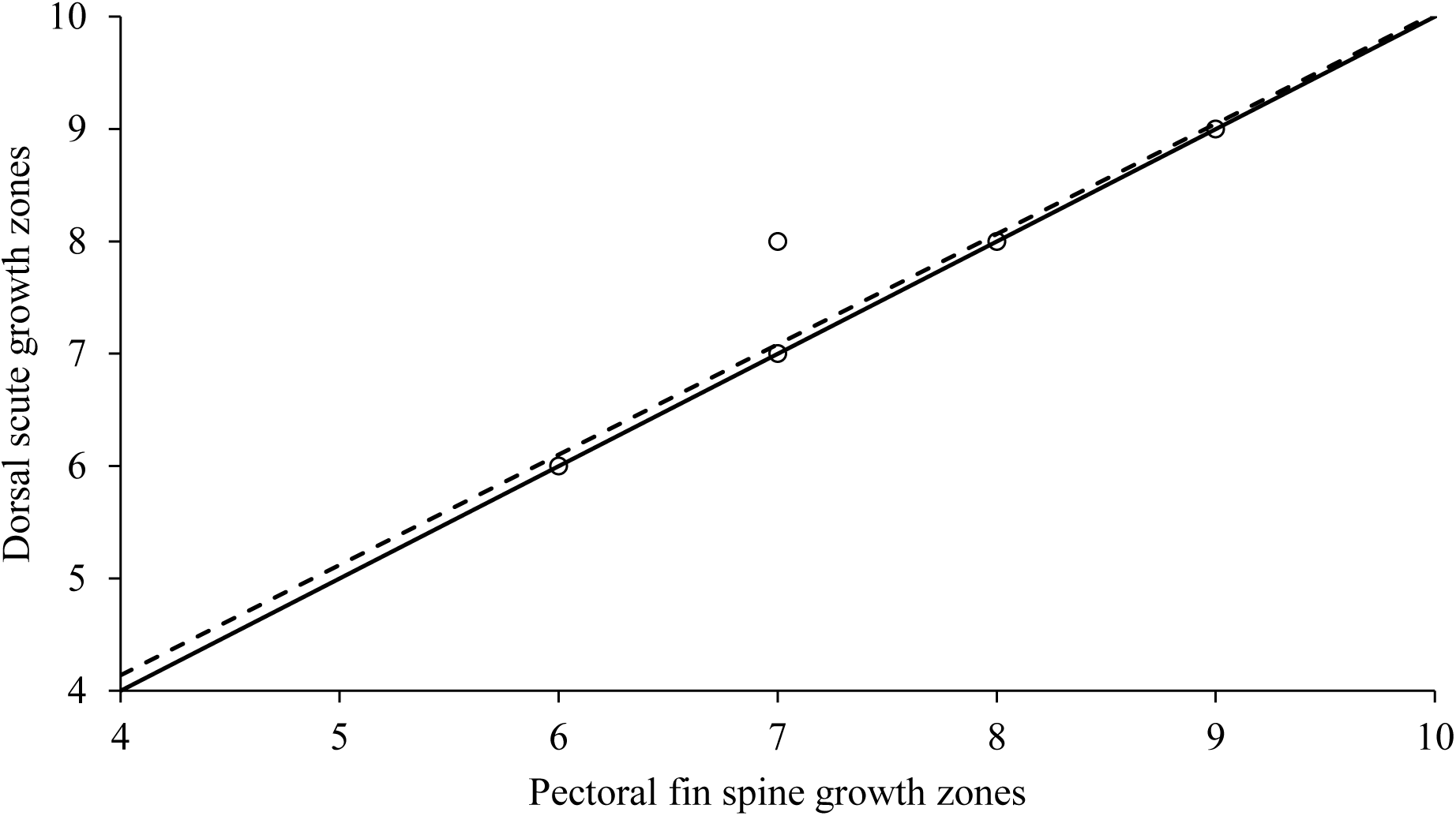
Regression plot of dorsal scute apical spine and pectoral fin spine growth zone counts for wild-caught Atlantic Sturgeon (n = 13; r^2^ = 0.95, *p* < 0.05). The dashed line represents the regression, and the solid line has a slope of one. The regression slope did not differ significantly from one (*p* = 0.336).

### Microchemistry analysis

Microchemical signatures across annuli from the same individuals were largely consistent between dorsal scutes and pectoral fin spines (e.g., Figure 5). Observed trends in these signatures showing a progressive decrease in Ba:Ca and an increase in Sr:Ca indicated movement across a salinity gradient from freshwater to the sea. While greater profile variability and magnitude in Sr:Ca and Ba:Ca ratios were observed in pectoral fin spine transects, overall correspondence in elemental patterns was observed across both structures. This was likely an artifact of the differences in overall transect length between structures—i.e., dorsal scutes were generally smaller than pectoral fin spines and had relatively compressed annuli that resulted in lower resolutions because less material was available to be ablated for chemical analysis.

**Figure 5.**
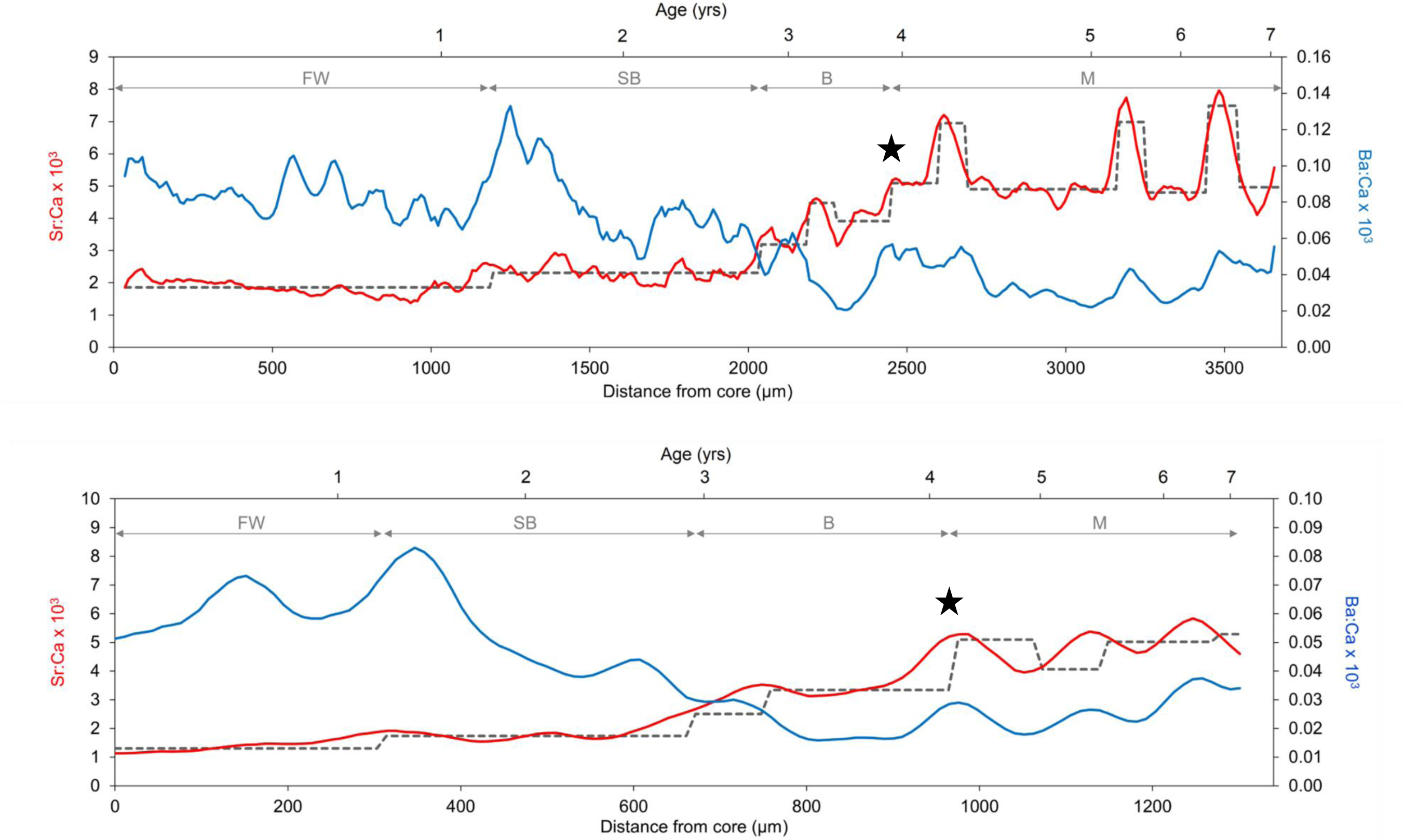
Pectoral fin spine (top panel) and dorsal scute apical spine (bottom panel) elemental profiles from a wild-caught Atlantic Sturgeon (USFWS ID-355160038). Derived Sr:Ca (red lines) and Ba:Ca (blue lines) are shown as five-point moving averages across transects that extend chronologically from the natal origin to the outer edge of the structure (see Figure 3). Dashed lines represent the mean Sr:Ca ratio observed in each regime (Rodionov, 2004). Estimated age and corresponding habitat delineations are provided on the upper axes (FW = freshwater; SB = seasonally brackish; B = brackish; M = marine; see Figure 1). The initial marine transition is indicated by a star, and assumed reentries into riverine habitat during the marine phase (i.e., simultaneous spikes in Sr:Ca and Ba:Ca signatures) are noticeable in both structures. Note, transect lengths vary based on structure size.

### Life history transitions and validation

Delineation of habitat use inferred from the microchemistry analysis of dorsal scutes and pectoral spines matched the expectation of an anadromous life-history, with habitat shifts in Sr:Ca indicating initial freshwater residence, followed by movements into progressively higher-salinity waters and eventual transition into fully marine habitat (Figure 6). The inferred initial marine transition generally coincided with the observation of a relatively large growth increment in the sectioned sample (Figure 3). Age estimates for the timing of this transition ranged from two to five years, with a mean age of 3.3 years (SD = 0.85; Table 1). Simultaneous peaks in Sr:Ca and Ba:Ca signatures that followed the initial marine migration were likely indicative of seasonal (i.e., within-year) movements back to estuarine habitat.

**Figure 6.**
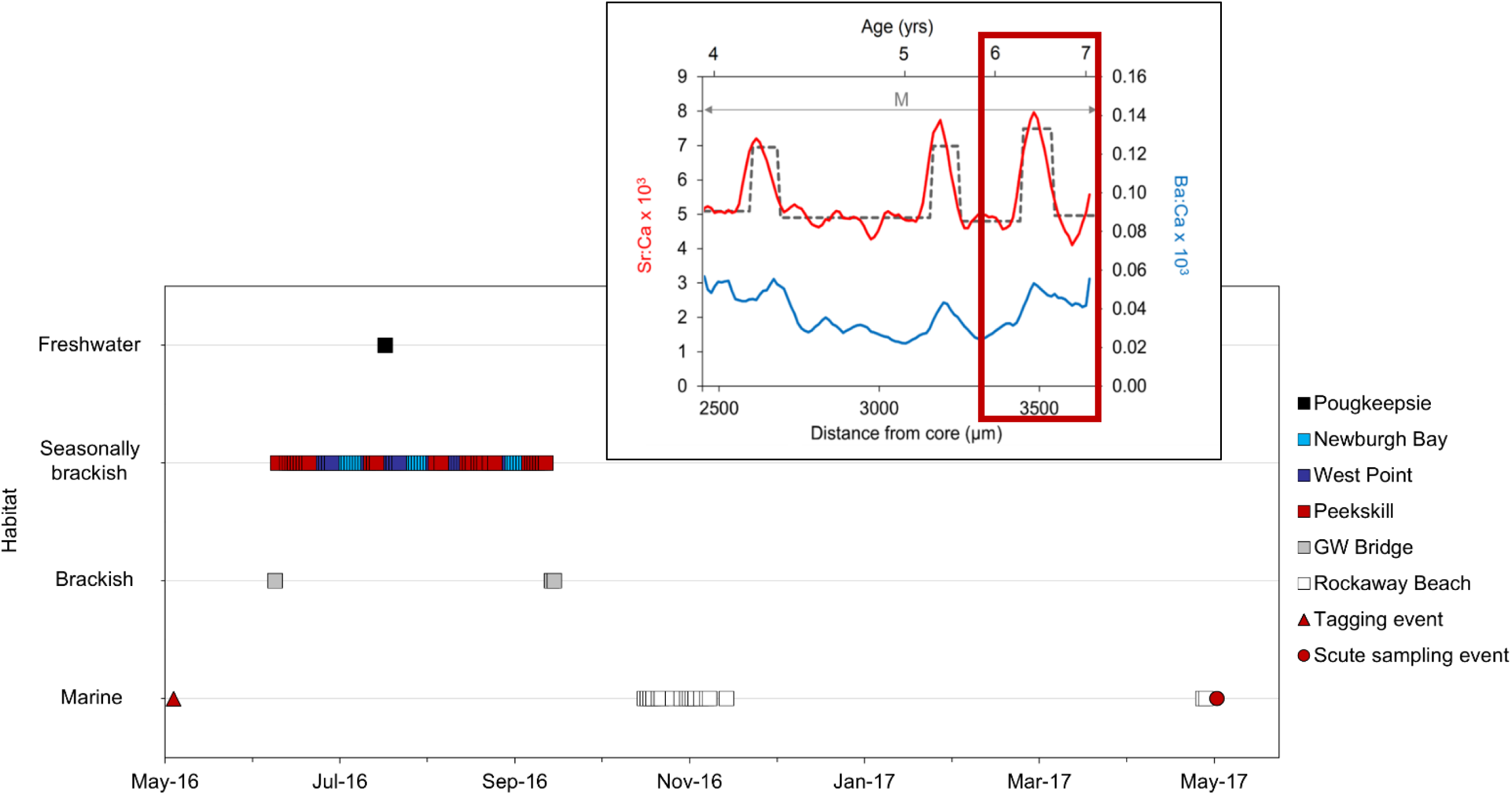
Comparison of acoustic telemetry detections showing habitat use of an Atlantic Sturgeon from the date of initial transmitter implantation until recapture and hard-part sampling (USFWS ID-355160038; May 4, 2016–May 9, 2017). Telemetry detections within the lower Hudson River, New York, and surrounding waters (see Figure 1) were used to validate inferences of habitat use from microchemical signatures of Sr:Ca and Ba:Ca in the final annuli (red box; ∼3400 μm; see Figure 3).

Detection history from acoustic telemetry monitoring was available for a single Atlantic Sturgeon that had been previously tagged with a unique-coded acoustic transmitter in marine waters off the Rockaways (VEMCO [Halifax, Nova Scotia]; V16-4H, 69 kHz; tag delay 70–150 s; estimated battery life = 3,650 d; Table 1). This individual female fish was from the Hudson River and was at-large from May 4, 2016, until capture and sampling on May 9, 2017, resulting in 7220 total detections, including 1462 marine detections and 5,758 riverine detections (Figure 6). During this time, the fish was detected migrating from the marine tagging location into the Hudson River, where it moved through brackish waters (18–30 ppt) and into seasonally brackish waters (0.5–18 ppt)—remaining near the salt-wedge from June through October, other than briefly entering freshwater habitat (< 0.5 ppt) in July—before leaving the river and returning to marine waters in November (Figures 1 and 6). These known locations were used to verify the habitat inferences from microchemistry analysis of the individual’s final annuli in both structures, which putatively corresponded to the period while the fish was at large (Figures 5 and 6). While short-term, seasonal migrations were more obvious from the pectoral fin spine, stage-specific transitions were readily identified from both structures (Figure 5).

## Discussion

Our study is the first to validate microchemistry inferences of environmental exposure for a wild-caught population of fish. The comparison of concurrent data from trace element patterns in dorsal scutes and pectoral fin spines with spatially referenced telemetry detections allowed for simple and categorical verification of habitat use inferences from microchemistry analysis. Demonstrable congruence between age estimates and trace-element ratio patterns between structures taken from the same fish indicates that dorsal scutes are unbiased timekeepers and contain a reliable chemical record of past environmental exposure. The ability to further delineate age-specific transitions between freshwater and marine environments from regime shifts in dorsal scute elemental profiles across annuli has broad management applications for Atlantic Sturgeon—particularly for data poor life-stages that may be difficult or impractical to study using traditional methods.

This study provides new information on Atlantic Sturgeon regarding the timing of ontogenetic transitions between riverine and marine habitats. Fish that were identified to the Hudson River (n = 11) remained in their natal system for two to four years of life before migrating to the sea. The timing of this initial transition into fully marine waters largely corresponded with Bain’s (1997) description of the ‘intermediate juvenile’ stage in the Hudson River, which is the transitional phase between ages three and six when fish gradually emigrate from the river during a period of rapid growth (based on length-frequency distributions of year class). The ability to accurately identify the age at which this transition occurs is essential to inform quantitative assessments for Atlantic Sturgeon populations. Because of the complex migratory history Atlantic Sturgeon, recent mark-recapture abundance estimates have largely focused sampling efforts on river resident fish that have yet to emigrate from their natal system to ensure that the population being sampled is closed (e.g. Schueller and Peterson, 2010; Baker et al., 2023). However, the assumption of closure has been difficult to validate without an understanding of the timing of ontogenetic transitions (Fox and Peterson, 2019). Inferences from dorsal scute microchemistry provide further support for these assumptions of closure and can be used to reliably define the ages at which specific cohorts first emigrate from their natal river system.

Major ontogenetic shifts in the microchemistry signatures from both structures suggest that the methodology was able to appropriately identify the timing and frequency of critical life history transitions for Atlantic Sturgeon. These inferences provide new data on Atlantic Sturgeon movements—both within and between management boundaries—that are necessary to identify age-specific threats associated that may be exacerbated by increased interactions with developed coastal landscapes throughout ontogenetic development. Furthermore, this information can be used to better define critical habitats outside of currently designated river systems (USOFR, 2017). Changes in Ba:Ca profiles in both structures during early river residence matched the expected signal for freshwater exposure, while observed regime shifts in Sr:Ca were consistent with downstream movements into progressively saline riverine habitats and subsequent residence in dynamic estuarine environments. Following the initial transition into marine waters, concurrent peaks in elemental profiles Sr:Ca and Ba:Ca profiles were indicative of large juveniles and sub-adults returning to the lower sections of coastal rivers during the warm, summer months. While the occurrence of non-spawning, marine-resident life stages of Atlantic Sturgeon in natal and non-natal river systems previously been reported from sampling efforts and tag recaptures (Dovel and Berggren, 1983; Van Eenennaam, 1996; Bain 1997), the relatively frequency of these migratory behaviors was unexpected.

This study demonstrated the strengths as well as potential limitations associated with both structures. Transect sizes of dorsal scutes were approximately three-times smaller than those of pectoral fin spines, while providing similar estimates of age and the timing of critical life-history transitions. Because LA-ICP-MS instrument time (and subsequent costs) are dependent on transect size, dorsal scutes are more practical for batch microchemistry analysis and may be preferred depending on desired outcomes and budgetary considerations. Age estimation from annuli counts was also relatively straightforward for dorsal scutes because of the lack of false annuli. Conversely, short-term, seasonal movements observed during the marine-resident stage were more obvious in pectoral fin spines. This suggests that larger transect size increases fine-scale resolution across each annulus by providing more ablated material for chemical analysis. As such, the decision of which structure to use is largely dependent on research goals. However, we suggest that the collection of both structures is warranted when sampling wild Atlantic Sturgeon populations. Although dorsal scute sampling is not currently authorized by NMFS without special dispensation, our methods for collection were shown to be minimally invasive for the life-stages we investigated (i.e, marine migrants > 500 mm FL) and could be easily incorporated into existing sampling protocols. As calcified structures, dorsal scutes are relatively inert and can be stored indefinitely for later analysis or potential submission to a centralized research repository.

The collection of dorsal scutes can complement ongoing research efforts for Atlantic Sturgeon by providing an additional tool to better define how individuals and populations are spatially dispersed across habitats. Inferences from microchemistry analysis provide critical data points on migrations and habitat use beyond those available from telemetry detections alone, including details on broadscale movements and habitat use that occur prior to sampling encounters or outside of monitored areas. Future work should focus on and refining the inferences from microchemistry analysis by investigating the kinetic relationship in Sr uptake in brackish waters as well as expanding the scope of analysis to include additional trace element profiles that can act as proxies for hypoxia (i.e., Mn:Ca; Jiang et al., 2022) and metabolism (i.e., Mg:Ca; Martin and Thorrold, 2005) or as potential indicators of spawning (i.e., Zn:Ca; Sturrock et al., 2015). While permitting considerations for endangered Atlantic Sturgeon currently limit the number of dorsal scutes that can be collected, expansion of this methodology to include fish from additional river populations is needed to investigate of clinal trends in the timing of critical life stage transitions and identify appropriate management strategies across the species’ coastwide range.

## Acknowledgements

Funding for this project was provided in part by the New York State Department of Environmental Conservation. Additional support for E. Ingram’s graduate studies was provided through the Pikitch Family Endowed Student Research Award and the Hudson River Foundation Mark B. Bain Graduate Fellowship. Initial methodology for dorsal scute sampling was determined based on early conversations with M. Kinnison, K. Lachapelle, and G. Zydlewski (University of Maine). The final protocol was refined on hatchery-raised fish at the Northeast Fishery Center in Lamar, PA, with support from S. Davis, T. Kehler, and M. Bartron (US Fish and Wildlife Service). Amendments to endangered species permitting were facilitated by M. Mohead and E. Markin (National Marine Fisheries Service). Laser ablation-inductively coupled plasma-mass spectrometry was completed with the assistance of D. Driscoll (SUNY College of Environmental Science and Forestry). We appreciate the invaluable field assistance provided by J. Zacharias, Captain C. Harter, and the crew of the RV Seawolf (Stony Brook University).

## Supporting Information

**Figure S1.**
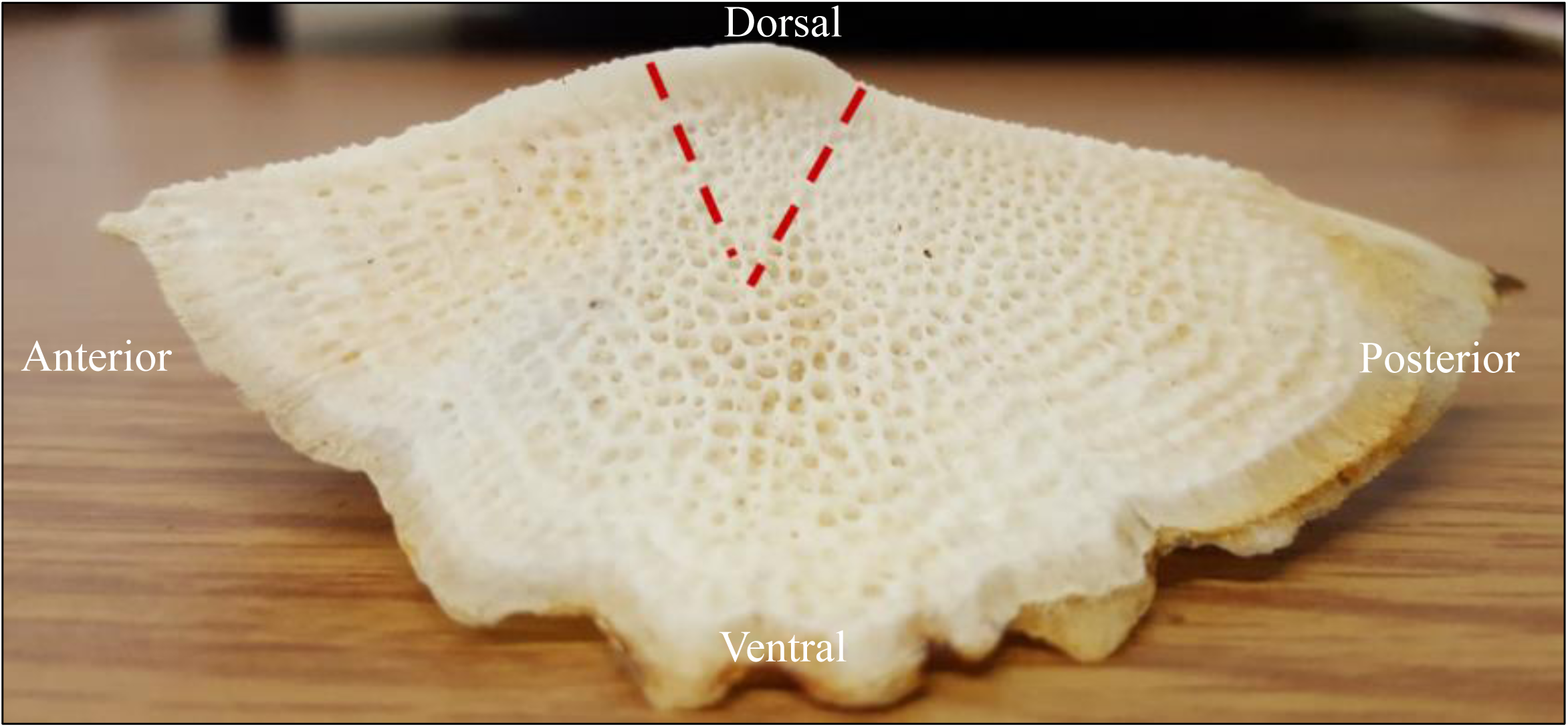
Image of Atlantic Sturgeon dorsal scute with sampling guidelines (dashed red lines). For aging and microchemistry analysis of wild-caught fish, a fine-toothed hacksaw was used to remove a wedge-shaped sample of the dorsal scute apical spine from the most-intact dorsal scute with two shallow, angled cuts perpendicular to the long axis of the fish.

**Figure S2.**
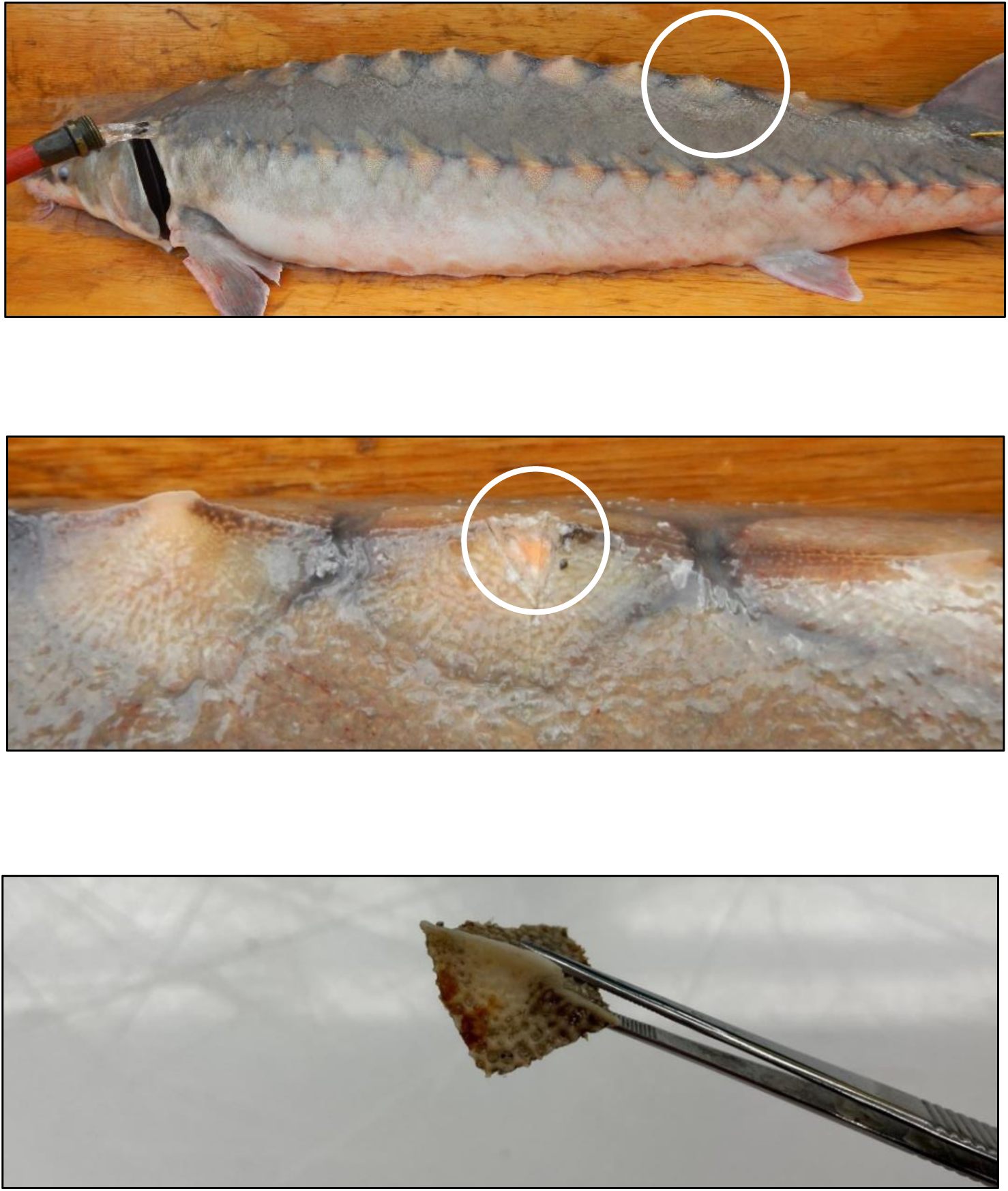
Atlantic Sturgeon captured in marine waters off the Rockaway Peninsula, New York, with the scute apical spine sampled from the 10^th^ dorsal scute (white circle).

## Appendix S1.

Detailed methods for the preparation of hard-part samples for aging and microchemistry analysis. This appendix supplements the ‘*Fish sampling*’ section of the main text methods

### Sample Preparation

Molds and epoxy bases—Two different-sized silicone ice cube trays (rectangular wells with dimensions of 2.0 x 5.0 x 1.5 cm and square wells with dimensions of 2.5 x 2.5 x 2.5 cm) were used as molds to accommodate variability in dorsal scute height and width. In the tray with square wells, the separating walls were removed with a razor blade to create rectangular wells that were slightly deeper than those in the other trays. Each well was filled with a thin layer of epoxy using Buehler EpoxiCure 2 Epoxy Resin (Lake Bluff IL, USA) and Buehler EpoxiCure 2 Epoxy Hardener (Lake Bluff IL, USA). The epoxy resin and hardener were weighed and mixed by volume according to the manufacturer’s directions. The epoxy was gently stirred for two minutes using a wooden popsicle stick to create a homogenous mixture and to remove any large air bubbles. The mixture was then poured into the silicone ice cube trays to completely coat the bottom of the wells, creating a base for the dorsal scute apical spine samples (i.e., dorsal scutes). Long slips of Rite in the Rain white all-weather paper (J. L. Darling Corporation, Tacoma, Washington, USA) were inserted into the wet epoxy mixture, leaving 3–4 cm of paper exposed and overhanging the wells to be used for subsequent labeling. The silicone trays with freshly poured epoxy bases were then repeatedly tapped on the lab bench 5–10 times to remove any remaining air bubbles and left to set for 24 hours.

Embedding dorsal scute samples—Once epoxy bases had hardened, dorsal scute samples were assigned to the wells depending on their width and height. Apical spines were placed in the silicone molds in an upright “A” position so that when they were later sectioned, the saw would make a transverse cut down the apex of the dorsal scute. Up to two samples could be placed into a single well with sufficient room to separate and section them individually. The samples were affixed to the epoxy bases using 3 –5 drops of liquid superglue (Loctite 0.35 oz, 10 g; Henkel, Rocky Hill, CT) per side. Each sample was manually held in place until it was able to remain upright on its own, typically until the superglue started to dry and become tacky. Each sample was labeled according to its unique dart identification number sequence. Once the samples were secured and the glue had dried, the silicone molds were filled with enough epoxy to completely cover the samples. Care was taken to ensure that epoxy filled in the area underneath the sample as well. Once filled, the wells were tapped on the lab bench to remove any remaining bubbles and left to dry for 24 hours.

Embedding pectoral fin spine samples—Pectoral fin spine samples (i.e., fin spines) were individually placed into Fisherbrand 2.0 mL MCT graduated natural microcentrifuge tubes (Fisher Scientific) and covered in epoxy (see above section). Tubes were then tapped on the lab bench approximately 5–10 times to remove as many air bubbles from the epoxy as possible. Tubes were left open to dry in an upright position for 24 hours. Once dry, each tube was cut open down the side using a razor blade to separate it from the epoxy-embedded fin spine. This resulted in a a cylindrical piece of epoxy containing the fin spine that was ready for sectioning.

### Sample Sectioning

Dorsal scute samples—Prior to sectioning the dorsal scute samples, each epoxy embedded sample was marked three times with a permanent marker. The first mark indicated the position of the initial cut needed to separate samples (if multiple samples were embedded in a single well). The second and third marks denoted the saw position needed to cut a thin section of the sample containing the entire profile of the “A-framed” sample from top to bottom. Samples were sectioned using a Buehler IsoMet low-speed sectioning saw (Lake Bluff IL, USA). The saw blade was dressed prior to sectioning and was set to revolve continuously at speed 7. The coolant reservoir of the saw was filled with deionized water while cutting. A 26.3g weight was added to the sectioning arm to encourage the saw to progress down through the dorsal scute. To prepare the dorsal scute sections for polishing, a petrographic microscope slide (56 x 27 x 1.2 mm) was labeled with the respective sample identification using a glass etching pen. The labeled petrographic slides were placed on a hotplate and pieces of Crystalbond 509 (Electron Microscopy Sciences, Hatfield PA) were used to affix each sampled section to the slide. The slide and sample were then allowed to cool completely prior to polishing.

Fin spine samples—Each cylinder of epoxy that contained a fin spine sample had excess epoxy removed from the before the final sectioning. Because of the length of each sample, multiple sections were cut to ensure that the entire profile of each sample was captured. Two weighted discs (76.6 g and 26.3 g) were added to the sectioning arm of the saw to encourage the blade to cut through each sample. Each sectioned sample was then attached to a petrographic microscope slide following the same procedure as for the dorsal scutes (see above section).

### Slide Preparation

Section mounting—The glass from an 8 x 10 picture frame was attached to a lab bench using 3M Scotch blue painters’ tape with Edge Lock #2093EL (St. Paul, MN) to secure all four corners and create a flat surface for sample polishing. Once the glass was affixed to the lab bench, 8” aluminum oxide abrasive film disks (3 μm and 30 μm) were cut into quarter segments. The adhesive backing from a 3 μm and 30 μm quarter segment was removed, and the film was then attached to the glass panel. The aluminum oxide abrasive quarters were saturated with deionized water to create a damp surface for sample polishing. A Kimwipe (Kimtech) barrier was created between the abrasive disks to reduce the amount of wastewater flow between grits. Suction cups from aquarium thermometers were wet and attached to the back of the petrographic slides to help create a firm hold on the slides when polishing.

Section polishing—The more abrasive 30 μm film was initially used to increase the rate of material removal and to achieve clarity and visibility of the annuli. Samples were polished in a figure-eight motion to ensure uniformity of the polished surface and to prevent the creation of sloped sections. The samples were regularly dried with Kimwipes and checked under a microscope to ensure they were not over polished, which could damage the samples and prevent accurate reading of the annuli. Once annuli were clearly visible, 3 μm abrasive film was used to increase clarity at a fine scale. This process was repeated for each dorsal scute and fin spine.

Final slide mounting—Once polished, individual dorsal scute sections were heated and removed from their slides and transferred to a newly labeled slide holding multiple samples. The dorsal scute sections were re-attached using Crystalbond 509 before removing the slide from the hotplate to cool completely. These multi-section slides retained a 3-mm border on all sides to allow them to be inserted into the laser ablation inductively coupled plasma mass spectrometry sample holder. This same method was repeated for the fin spine samples. Once all slides were cooled, they were stored in a Petrographic slide box.

